# Neural substrates of working memory updating

**DOI:** 10.1101/853630

**Authors:** Gal Nir-Cohen, Yoav Kessler, Tobias Egner

**Affiliations:** Department of Cognitive and Brain Sciences, Ben-Gurion University of the Negev; Department of Psychology, Ben-Gurion University of the Negev; Zlotowski Center for Neuroscience, Ben-Gurion University of the Negev; Center for Cognitive Neuroscience, Duke University; Department of Psychology and Neuroscience, Duke University

## Abstract

Working memory (WM) needs to protect current content from interference and simultaneously be amenable to rapid updating with newly relevant information. An influential model suggests these opposing requirements are met via a basal ganglia (BG) - thalamus gating mechanism that allows for selective updating of prefrontal cortex (PFC) WM representations. A large neuroimaging literature supports the general involvement of the PFC, BG, and thalamus, as well as posterior parietal cortex (PPC), in WM. However, the specific functional contributions of these regions to key sub-processes of WM updating, namely gate-opening, content substitution, and gate closing, are still unknown, as common WM tasks conflate these processes. We therefore combined functional MRI with the reference-back task, specifically designed to tease apart these sub-processes. Participants compared externally presented face stimuli to a reference face held in WM, while alternating between updating and maintaining this reference, resulting in opening vs. closing the gate to WM. Gate opening and substitution processes were associated with strong BG, thalamic and fronto-parietal activation, but – intriguingly - the same activity profile was observed for sensory cortex supporting task stimulus processing (i.e., the fusiform face area). In contrast, gate closing was not reliably associated with any of these regions. These findings provide new support for the involvement of the BG in gate opening as suggested by the gating model, but qualify the model’s assumptions by demonstrating that gate closing does not seem to depend on the BG, and that gate opening also involves task-relevant sensory cortex.

## Introduction

Working memory (WM) refers to our ability to temporarily hold information in mind, manipulate it, and update it in the service of goal-directed behavior (Cowan, 2017; Oberauer et al., 2018). Models of WM have long emphasized the tension between its maintenance and updating functions (Badre, 2012; Fallon et al., 2017; Frank et al., 2001; Miller & Cohen, 2001; O’Reilly, 2006): current WM content has to be shielded from interference by irrelevant information, while at the same time being amenable to updating when new goal-relevant information appears.

An influential neurocognitive theory addressing this dilemma is the PBWM (prefrontal cortex, basal ganglia, and WM) network model, which postulates a selective input-gate for WM (Frank et al., 2001; O’Reilly & Frank, 2006). Specifically, the model proposes a basal ganglia (BG) gating mechanism that separates perceptual input, represented in sensory cortex, from WM representations, maintained in (or via) dorsolateral prefrontal cortex (dlPFC). By default, the gate is closed, thus enabling robust maintenance/shielding of WM content. However, in response to salient signals, like task-relevant stimuli or reward cues, the BG gate opens (based on phasic dopaminergic input from the midbrain), allowing for the inflow of new information into WM via a thalamus-PFC pathway.

There is a vast neuroimaging literature supporting the assumption that the dlPFC – usually in conjunction with the medial prefrontal cortex (mPFC) and posterior parietal cortex (PPC) – contributes to the maintenance of WM content (e.g., (D’Esposito & Postle, 2015; Feredoes et al., 2011; Nee et al., 2013; Roth et al., 2006). The same broad set of regions, often referred to as the frontoparietal network (FPN), has also been implicated in updating WM, as inferred from N-back (e.g., Owen et al., 2005), AX-CPT (e.g., Lopez-Garcia et al., 2016) and task switching studies (e.g., Kim et al., 2012). These regions have therefore been referred as the core network of WM (Harding et al., 2015; Johnson et al., 2019; Rottschy et al., 2012). A smaller set of studies has also provided evidence to support the involvement of the BG (Chatham & Badre, 2015; Cools et al., 2007; Murty et al., 2011) and/or the midbrain (D’Ardenne et al., 2012; Murty et al., 2011) in WM updating. More specifically, the involvement of the BG in gating goal-relevant information into WM – a key sub-component of WM updating - has been implicated in several studies. For example, previous studies reported BG involvement during switching attention between objects (Cools et al., 2004; van Schouwenburg et al., 2014) and tasks (Leber et al., 2008) Moreover, Van Schouwenburg et al., (2010) found that the BG mediated the connectivity between the PFC and visual cortex during attentional shifts, triggered by a bottom-up cue. Finally, McNab & Klingberg, (2008) demonstrated that activity in both the PFC and BG preceded the selection of relevant information for WM maintenance, and that this activity was associated with individual differences in WM capacity, resonating with the notion that capacity is related to filtering (i.e., gating) ability (Vogel et al., 2005).

While there is broad agreement on the FPN, BG, and thalamus being the key players in WM input gating, the mapping of these regions to the *processes* underlying WM gating and updating is presently unclear. These processes include opening the gate to allow information into WM, modifying the relevant items while removing outdated information, and returning to a closed-gate, perceptually-shielded state when updating is complete. The reason for a lack of such process-specific brain mapping in the prior literature is mainly due to task impurity. For instance, standard N-back, AX-CPT, and task switching protocols conflate item encoding, updating, substitution, and other processes, and do not provide a means to differentiate gate opening, gate closing, substitution and item removal processes (discussed in Ecker et al., 2010; Rac-Lubashevsky and Kessler, 2016a, 2016b; Kessler et al., 2017; Lewis-Peacock et al., 2018).

The goal of the present study was therefore to examine potential functional specialization in the WM network with respect to above-described sub-processes involved in WM updating. To this end, we paired functional magnetic resonance imaging (fMRI) with the recently developed “reference-back” task (Rac-Lubashevsky & Kessler, 2016a, 2016b, 2018), which has been shown to successfully disentangle processing costs associated with four key WM updating operations: (1) opening the gate to WM, (2) updating information in WM – which may either take the form of reinforcing current content or (3) substituting old with new information, and (4) closing the gate in order to enable robust maintenance of the newly updated information. In addition, we examined the neural correlates of being in an “updating mode” (see Kessler & Oberauer, 2014). Unlike the processes described above, the updating mode refers to the *state* of the gate to WM– whether it is open for new input or not.

By interrogating neural responses in the BG, FPN, and thalamus, as well as in visual regions with known sensitivity to our task stimuli (see below), we observed distinct patterns of neural substrates supporting the different WM updating processes. Whereas dlPFC, BG, and thalamus were preferentially involved in the gate opening process, parietal cortex also contributed to this process, but additionally displayed a stronger contribution to substitution. In contrast, these regions were not involved in gate closing.

## Materials and Methods

### Participants

To mitigate the dangers of false-positive and –negative findings, we based our sample size on effect size estimation (Button et al., 2013). Specifically, a recent meta-analysis of a large fMRI data set indicated a moderate effect size for WM task contrasts (Poldrack et al., 2017), For a desired power of 0.8 to detect this size of effect in within-subjects contrasts, under assumption of a conservative (low) level of correlation between paired observations (r=0.3), we aimed for a minimal sample size of N=45 (based on GPOWER, Erdfelder et al., 1996).

61 healthy students from Ben-Gurion University of the Negev participated in the experiment in exchange for monetary compensation. 13 participants were excluded from the analysis due to technical problems with the MRI during the scan (6), extensive head movements (2) or a low accuracy rate (5; <80%). The final sample included 48 participants (29 females; age M=25.5, SD=2 years). All participants were right-handed and reported normal or corrected-to-normal vision. None of the participants had any history of neurological or psychiatric problems. The experiment was approved by the Helsinki committee of the Soroka Medical Center, Beer Sheva, Israel.

### Stimuli

The reference-back task used 8 face images of neutral facial expression (4 males, 4 female; two faces per block) from FEI faces database (http://fei.edu.br/~cet/facedatabase.html). The faces were displayed inside blue (RGB values: 0,0,255) and red (RGB values: 255,0,0) colored frames (see Figure 1A). The faces’ diameter was approximately 180 pixels (4.76 centimeters, subtending a visual angle of 2.7° from a 100 centimeters viewing distance). The frame’s dimensions were 380×380 pixels (10×10 centimeters), subtending a visual angle of 5.7°. The use of face stimuli (in combination with an independent “localizer” scan) enabled us to identify the fusiform area (FFA; Kanwisher et al., 1997) in order to assess neural task stimulus processing in visual cortex as a function of WM updating operations. The localizer task employed gray scale images of famous familiar faces, unfamiliar faces, buildings, objects and scrambled objects. Those images were presented within an elliptical shape (14.5×8 centimeters, 8.3°x4.5°) against a black background. All stimuli were projected on a screen at the back of the scanner bore, and viewed via a mirror affixed to the headcoil.

**Figure 1:**
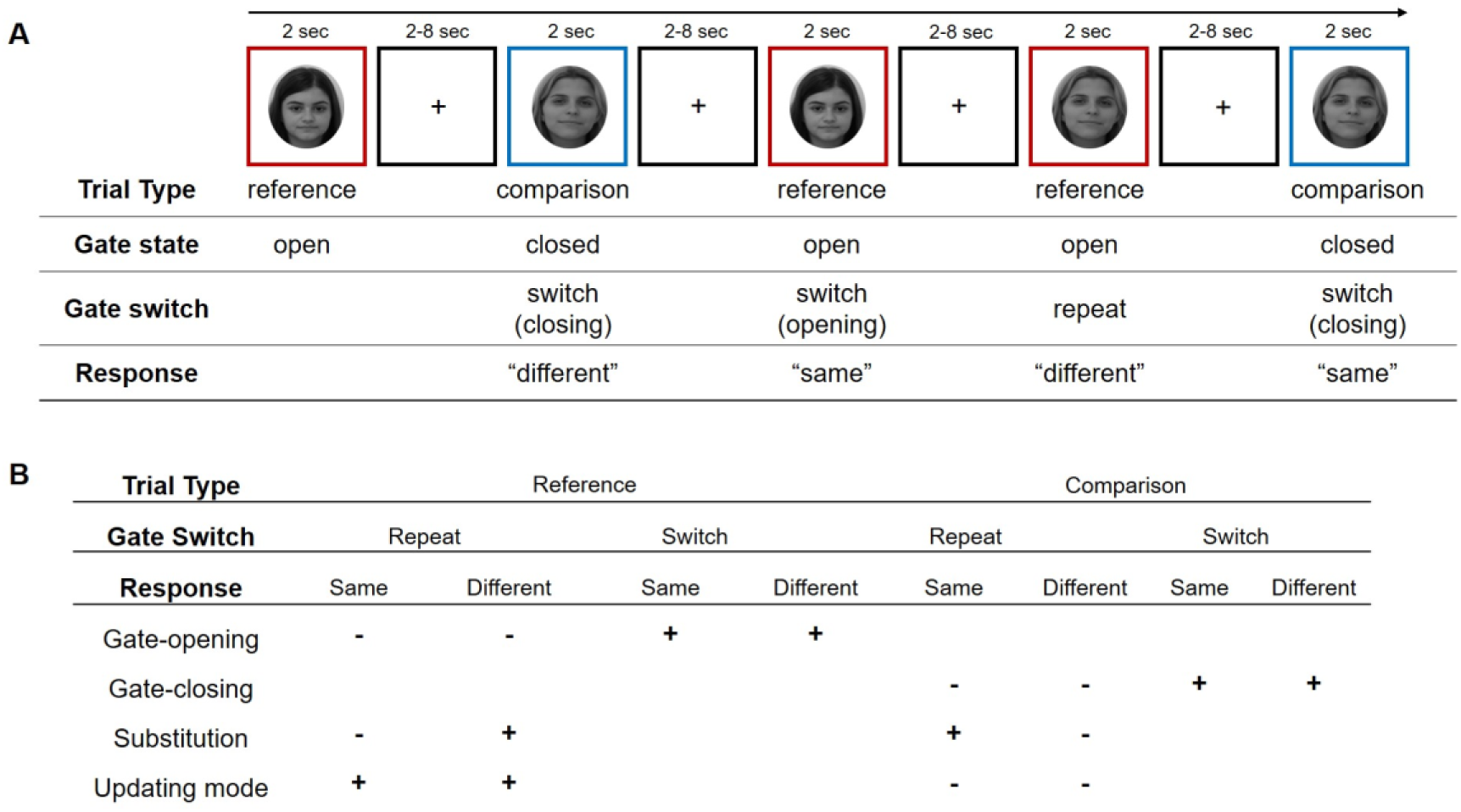
(A) : An example sequence of reference (red frame) and comparison (blue frame) trials in the reference-back task (top), along with an illustration of the putative states of the WM gating process and responses required (bottom). (B) Contrast weights for defining distinct WM updating sub-processes.

### Procedure

The scanning session took about an hour, in the following sequence: anatomical structural scan (10 min); 1-back task with face stimuli, serving as a face localizer task (10 min); and four 80-trials blocks of the reference-back task (30 min). The participants completed a behavioral practice session (2 blocks) of the reference-back task a day or two prior to the experimental session in the scanner.

### The reference-back task

We employed the reference-back task (Rac-Lubashevsky & Kessler, 2016a, 2016b), which enabled us to disentangle WM updating sub-processes (i.e., gate opening, gate closing, and substitution). This task is based on the N-back task. In the standard N-back task the participant is presented with a sequence of stimuli, and is asked to decide whether the current stimulus is identical or not to the stimulus presented N trials before (Jonides et al., 1997; Owen et al., 2005). Since updating, including various sub-processes, takes places in each trial of the n-back task, it is difficult to isolate the sub-process and identify their distinctive neural markers. In order to overcome this limitation, the reference-back paradigm was developed.

This task is composed of two trial types: reference and comparison. Specifically, in each trial (see Figure 1A), a face stimulus was presented inside a red or a blue frame, and the participant was required to indicate whether or not the stimulus was identical to the one presented in the *most recent red frame*. Trials involving a red frame are denoted *reference trials*. In these trials, the participant must first compare the presented stimulus to the one held in WM, i.e., the face that appeared in the previous red frame (making a same/different decision); the participant then has to update his/her WM with the stimulus that appears in the present trial, which serves as the reference for future trials. Trials involving a blue frame are denoted *comparison trials*. Like in red frame trials, the participant is required to make a same/different decision between the currently presented face and the reference held in WM; however, unlike in reference trials, WM does not have to be updated, because blue-framed faces do not serve as a reference for future trials.

Accordingly, both reference and comparison trials involve a same/different decision against a WM referent, but only the former require WM updating. This means that the gate to WM should be open in reference trials but kept closed in comparison trials. By considering the state of the gate on the previous trial, this protocol further allows one to distinguish between trials where the gate needs to be opened and trials where the gate needs to be closed. Specifically, trials in which the previous trial-type is repeated (e.g., two reference trials in a row) do not entail a change in the state of the gate: the gate remains open for successive reference trials and it remains closed for successive comparison trials. However, switching from a comparison trial to a reference trial requires gate opening, while switching from a reference to a comparison trial requires gate closing (see Figure 1 for a trial-by-trial example).

Consequently, the reference-back task enables one to distinguish among WM updating sub-processes using three pre-defined orthogonal contrasts. These contrasts were utilized in previous studies, showing robust behavioral effects (Kessler, 2017; Rac-Lubashevsky & Kessler, 2016a, 2016b) as well as EEG correlates (Rac-Lubashevsky & Kessler, 2018), and association with spontaneous eye blink rates, an index of central phasic dopaminergic activity (Rac-Lubashevsky et al., 2017). Moreover, reaction time costs for these contrasts demonstrate split-half reliabilities of .85-.86, and are correlated with performance in the standard N-back task (Rac-Lubashevsky & Kessler, 2016b).

The contrasts were defined as follows (also, see Figure 1B). (1) *Gate opening* is the difference between reference-switch and reference-repeat trials. All reference trials require the gate to WM to be open but only the former, where participants are switching from a comparison to a reference trial, involves the process of gate opening. Using a similar logic, (2) *Gate closing* was defined as the difference between comparison-switch and comparison-repeat trials – all comparison trials require a closed gate, but only on trials where participants switch from a reference to a comparison trial does the process of gate closing take place. Importantly, since each of the two face stimuli can appear in each of the conditions, the above contrasts are orthogonal to the correct response (being “same” or “difference”). Lastly, (3) *Substitution* refers to replacing old with new information in WM, which occurs on those reference trials where the current stimulus does not match the previous reference (see also Ecker et al., 2010). In order to substitute old with new information, the irrelevant previously-updated information should be removed (Ecker et al., 2014; Kessler, 2018; Lewis-Peacock et al., 2018). In the reference-back paradigm, removal and substitution are coupled (i.e., each time an item is substituted, the previous one should be removed; but see Kessler, 2018, for a version of this paradigm that enables observing the after-effects of removal by n-2 repetition costs). Hence, in the current study we are not attempting to isolate activity specifically related to removing items from memory, and part of the activations identified in the substitution contrast may reflect such a removal process.

It is important to de-confound substitution from the difference between making a “same” versus a “different” response, and this can be achieved by using the difference between “same” and “different” responses in comparison trials as a baseline. Accordingly, substitution is calculated as an interaction contrast, reflecting a larger difference between “same” and “different” responses in reference trials than in comparison trials: (“different” _reference_ – “same” _reference_) – (“different” _comparison_ – “same” _comparison_).

In addition, the reference-back design enabled us to examine the differential neural activity of being in an open-gate state (“update mode”, see Kessler and Oberauer, 2014, 2015) compared to a closed-gate state. Accordingly, the *updating mode* is defined by the overall difference between reference trials, where the WM referent has to be updated, and comparison trials, where the referent does not have to be updated. The updating mode contrast only involved trial-type repetition trials, in order not to confound it with gate-switching. Thus, while *substitution* refers to the situation where updating involves replacing of the old referent with a new one (and possibly also includes removing the now-irrelevant item, see Lewis-Peacock et al., 2018), the *updating mode* refers to the more general situation of being in an open-gate state, regardless of whether the referent has to be replaced (“different” trials) or not. These process designations, and the specific contrasts isolating the different updating operations are shown in Figure 1.

In mapping these reference-back gating costs onto the PBWM model, it should be noted that the latter assumes that the WM gate is closed by default and only opens transiently to relevant inputs, whereas the reference-back contrasts assume the gate to remain open following an updating event (a reference trial) until the next input is evaluated. The fact that robust gate-opening costs are reliably observed (Kessler & Oberauer, 2014, 2015; Rac-Lubashevsky et al., 2017; Rac-Lubashevsky & Kessler, 2016a, 2016b) argues against the PBWM model assumption, because such costs should not be obtained if the gate were closed automatically following each updating event (that is, reference trial RTs should not differ as a function of the preceding trial being a reference or comparison trial). However, we suggest that the PBWM model can be reconciled with these data via the plausible assumption of context-sensitive gating policies (e.g., Bhandari & Badre, 2018). To wit, in situations where updating is rarely required and distracters are frequent, it would make sense to keep the gate closed by default. By contrast, when updating is required frequently, as in the present task (on 50% of the trials), it would be more efficient to maintain the gate state from the previous trial until the next input is observed, since this policy would minimize the number of gate state switches (c.f. Kessler & Oberauer, 2014).

Each block of the reference-back task started with a reference trial, to which the participants did not respond. Then, in each subsequent trial, a framed face was presented for 2 seconds, followed by a blank inter-trial interval for 2, 4, 6 or 8 seconds. Each of the eight conditions (Trial-Type * Gate-Switching * Response) was presented 10 times in each block, resulting in a total of 320 trials (40 trials per conditions). The order of trials, as well as the duration of the inter-trial interval jitter, were determined using Optseq (Free-Surfer analysis tools; Greve, 2002). Each of the four experimental blocks involved two face stimuli from the same gender. The stimuli were changed from one block to another and were counterbalanced between participants. We employed different faces in each block to avoid contributions of long-term memory to performance. Moreover, using only two faces within a given block ensures that there is a high potential for interference, which promoted the use of WM over familiarity-based strategies (e.g. Szmalec et al., 2011). Note that the contrasts of interest (Fig. 1B) are orthogonal with respect to whether a specific face is repeated from one trial to the next, thus preventing face stimulus repetition suppression effects from confounding our results.

### FFA Localizer Task

As an FFA localizer task, we employed a block-design 1-back task (taken from Avidan et al., 2014). Different stimulus categories (familiar faces, unfamiliar faces, buildings, objects, and scrambled objects) were presented in 10 second blocks, with 6 seconds intervals between blocks. Within each block, ten images were presented, each for 800ms followed by 200ms inter-trial interval. Within each block, nine images were unique whereas one image was presented twice in a row. The participants were asked keep track of the stimuli, and to press a key each time an image was presented twice in a row (1-back). There were seven repetition of each block type.

### fMRI data acquisition and preprocessing

All fMRI data were collected at the Brain Imaging Research Center, Soroka Medical Center, Beer-Sheva, using a 3-T Philips ingenia MRI scanner. The scanning of each participant started with a 3D structural scan, acquired by a T1-weighted sequence that yielded high resolution images of 1^3^ mm voxel size with matrix of 256×256 for 170 slices. Functional data were collected by a T2*-weighted sequence (TR = 2000 msec, TE = 35 msec, flip angle = 90°). 35 slices were scanned in ascending order with a 96×96 matrix size, 2.61×2.61 mm voxel resolution with 3 mm thickness. A total of 215 volumes were acquired. Behavioral responses were recorded using a two-keys box the participants held in their right hand and pressed with their index or middle finger for different and same responses, respectively. Imaging data were preprocessed and analyzed using Statistical Parametric Mapping 12 (Welcome Trust Centre for Neuroimaging, London, UK; http://www.fil.ion.ucl.ac.uk/spm). Each participant’s functional images were re-aligned, co-registered to the anatomical image and slice-time corrected. Then, the images were normalized into MNI space (with a 2^3^ mm voxel size interpolation) and smoothed using a 6 mm gaussian kernel to full width at half maximum (FWHM).

### Statistical data analysis

In a 1^st^-level analysis, for each participant a task model was constructed with event-based stick functions, convolved with a canonical HRF, and high-pass filtered (128s) to remove low-frequency signal drift. The subject-level task matrices included one regressor for each of the eight conditions resulting from the 2 (trial type: reference vs. comparison) x 2 (gate switch: repeat vs. switch) x 2 (response: same vs. different) factorial design shown in Figure 1B. The models also included a regressor accounting for error trials, null trials, the grand mean, and six head-movements regressors. Four linear contrasts were defined to estimate activation for gate opening, gate closing, substitution, and updating mode, respectively, as explained in the task description above (See Figure 1B). For the FFA localizer task, the individual task models were constructed in the same manner, but coding for blocks of face stimuli vs. non-face stimuli, which were contrasted against each other.

The individual participants’ contrast images were then submitted to a 2^nd^-level one-sample *t*-test group-level analysis, where participants were treated as random effects. We pursued two broad sets of fMRI analyses, the first being an exploratory whole-brain analysis, and the second being a region-of-interest (ROI) analysis grounded in our *a priori* interest of closely interrogating the role of different nodes of the WM network (the constituent parts of the FPN, the BG nuclei, and the thalamus) and the FFA in distinct aspects of the WM updating process. Accordingly, we took a very conservative approach for guarding against false-positives in the exploratory analysis, using a voxel-based family wise error (FWE) with a threshold of *p* < 0.05 and a minimum cluster size (*K*_*E*_) of 10 significant voxels for the whole-brain analysis, and employed a less conservative approach for the *a priori* ROI analysis (which in turn is less likely to avoid false-negatives) by using a voxel-based false discovery rate (FDR) with a threshold of *p* < 0.05. Note that we used voxel-based rather than cluster-based thresholding throughout to bypass recent concerns about common cluster-based correction approaches (Cox et al., 2017; Eklund et al., 2016). Based on a large WM literature (D’Esposito & Postle, 2015; Frank et al., 2001; Harding et al., 2015; McNab & Klingberg, 2008; Nee et al., 2013; O’Reilly & Frank, 2006; Rottschy et al., 2012), the WM network ROIs were defined anatomically, using the WFU PickAtlas toolbox (Maldjian et al., 2003) masks of the BG and thalamus (using AAL labels; Tzourio-Mazoyer et al., 2002), and assembling an FPN mask, using Brodmann areas, by combining masks of the dorsolateral PFC (BA8, BA9, BA46), medial PFC/ACC (BA24, BA32), and the posterior parietal cortex (BA7, BA40). We specified a functional ROI of the FFA via the localizer scan. Finally, in addition to searching for significant clusters of activation within each ROI, we also ran the above-mentioned contrasts on mean activity (beta estimates) extracted from each ROI. Bayes factors favoring the alternative (BF_10_) and the null (BF_01_) hypotheses and were calculated using JASP software (JASP Team, 2019) with a default prior. The latter is especially important in order to establish meaningful null effects.

## Results

### Behavioral results

All the conditions, along with the four *a priori* contrasts of interest were tested on both response time (RT) and accuracy; mean RTs for the key conditions are shown in Figure 2, and descriptive and inferential statistics are presented in Tables 1 and 2, respectively. The results fully replicated those of the original studies on the reference-back paradigm (Rac-Lubashevsky & Kessler, 2016a, 2016b): As can be seen in Figure 2, mean RT for reference trials was substantially slower than for comparison trials, reflecting the cost of the WM updating mode (54ms, *p* < 0.001). As shown in Figure 2A, switch trials were found to be significantly slower than repeat trials, both in reference trials and in comparison trials, reflecting the costs of gate-opening (72ms, *p* < 0.001) and gate-closing (52ms, *p* < 0.001), respectively. Finally, the interaction contrast comparing “same” and “different” response conditions between reference and comparison trials revealed a robust substitution cost (92ms, *p* < 0.001). In summary, the behavioral results showed that our adaptation of the reference-back protocol was successful in revealing the behavioral signatures of updating, gate opening and closing, and substitution processes in WM, thus providing a solid basis for interrogating the fMRI data for neural substrates of these processes.

**Table 1:** Descriptive statistics of the RT and accuracy data.

| Conditions |  | Response Time |  |  | Accuracy |  |
| --- | --- | --- | --- | --- | --- | --- |
| Trial Type | Gate switch | Response | Mean (ms) | SD | Mean (%) | SD |
| Reference | Repeat | Same | 694 | 113 | 98 | 3.02 |
|  |  | Different | 882 | 158 | 97 | 3.28 |
|  | Switch | Same | 773 | 160 | 98 | 2.62 |
|  |  | Different | 945 | 199 | 96 | 4.48 |
| Comparison | Repeat | Same | 688 | 116 | 98 | 2.47 |
|  |  | Different | 782 | 145 | 98 | 3.15 |
|  | Switch | Same | 716 | 114 | 98 | 2.93 |
|  |  | Different | 857 | 165 | 97 | 2.67 |

**Table 2:**
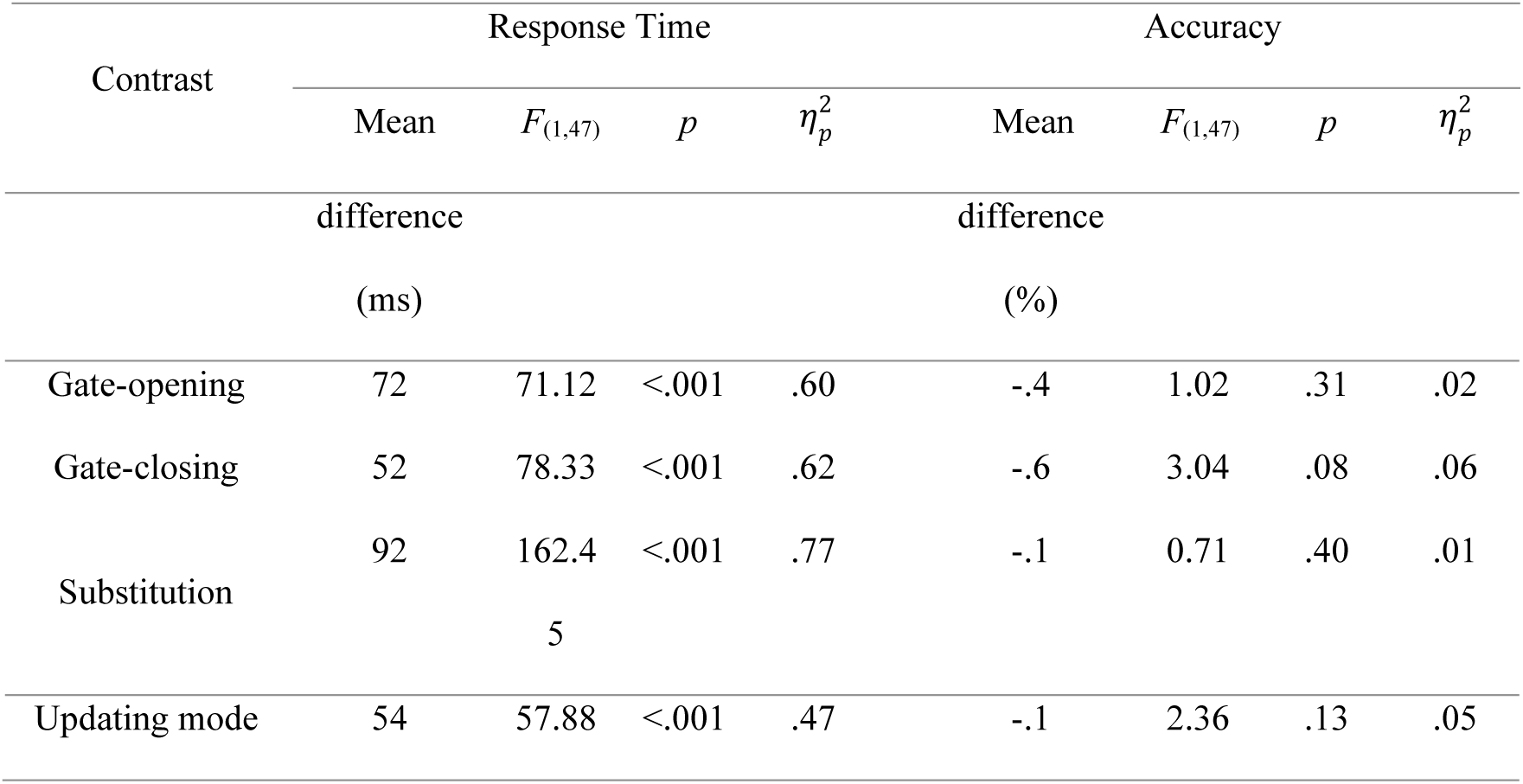
Summary of the inferential statistics of the RT and accuracy data analysis

**Figure 2:**
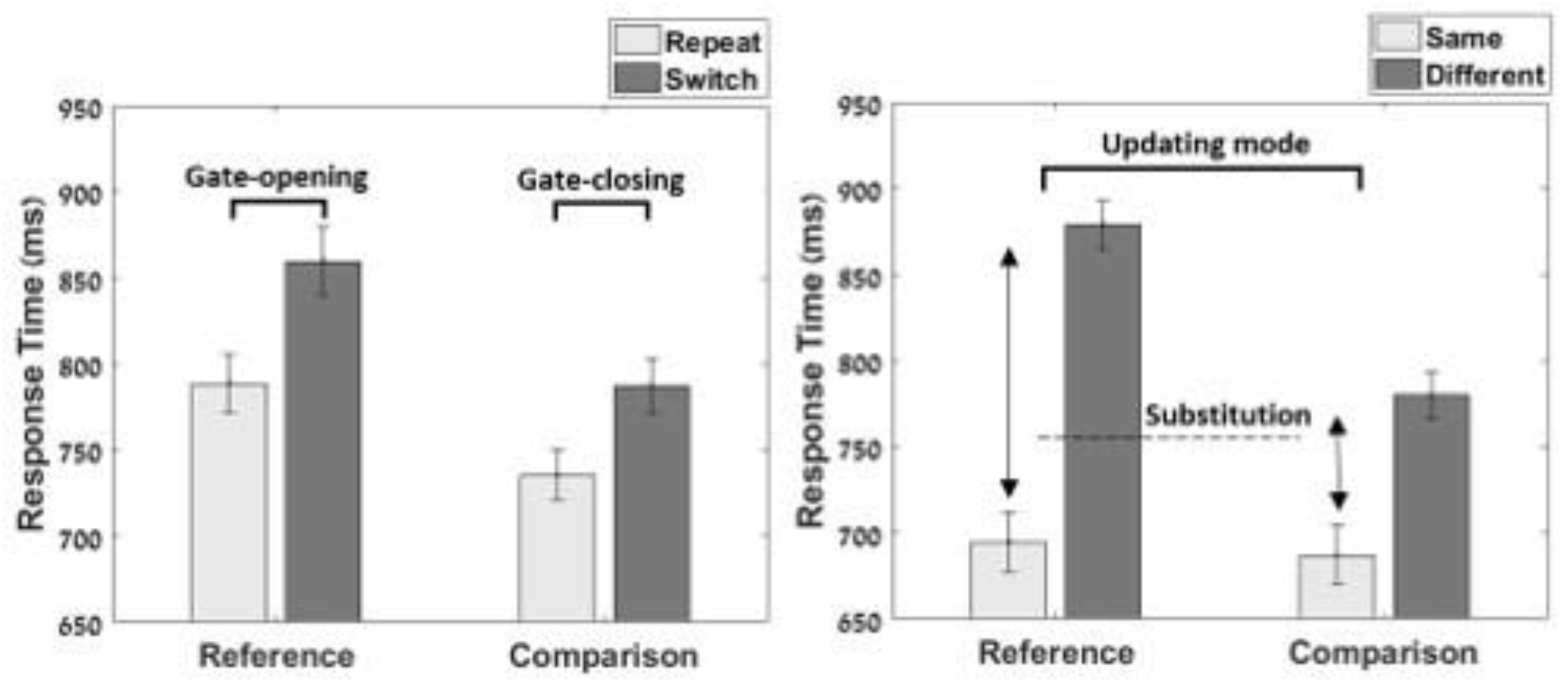
Mean response time (and standard error of the mean) for reference and comparison trials are displayed (A) as a function of whether the condition was repeated or switched to, and (B) of whether the stimulus/response was the same or different to the reference stimulus held in WM. The figure also highlights the four key contrasts defining gate-opening, gate-closing, substitution and updating mode.

### Imaging results

#### Exploratory Whole-brain analysis

We began with an exploratory whole brain analysis of each contrast of interest using a conservative correction threshold (voxel-wise FWE *p* < 0.05, *K*_*E*_ > 10). Dorsal views of cortical activations revealed by each contrast are presented in Figure 3 (for full list of activated clusters, see Table 3). The process of *gate opening* (required in reference trials that follow a comparison trial) was associated with increased activation in dorsal and dorsomedial frontal and parietal regions, with particularly large clusters of activity observed in the posterior and medial aspects of the PPC, including the precuneus. Additionally, gate-opening was associated with activity in the thalamus, as well as an extensive posterior cluster stretching from the cuneus into parts of visual cortex, including the fusiform gyrus (for full list of activated clusters, see Table 3). Given that this contrasts controls for basic visual input (which is equated between reference and comparison trials), the latter data suggest that the process of gating visual information into WM may be directly reflected in enhanced activity in relevant visual regions (see also FFA ROI-based analyses, below).

**Table 3:** Whole-brain analysis’ list of peak activation in MNI coordinates; Z refer to z-score at peak activated voxel

| Region | HM | $K_E$ | MNI coordinates | | | Z |
| --- | --- | --- | --- | --- | --- | --- |
|  |  |  | x | y | z |  |
| Gate opening (reference switch > reference repeat) |  |  |  |  |  |  |
| Precuneus |  | 4,647 | 0 | -70 | 34 | 7.33 |
| Fusiform gyrus | R | 133 | 30 | -74 | -10 | 7.05 |
| SMA | L | 43 | -2 | -8 | 52 | 5.56 |
| MFG | L | 20 | -22 | 14 | 58 | 5.41 |
| IFG | L | 37 | -50 | 24 | 22 | 5.39 |
| Thalamus | L | 18 | -8 | -18 | 10 | 5.34 |
| Insula | L | 17 | -28 | 28 | 0 | 5.25 |
| Gate closing (comparison switch > comparison repeat) |  |  |  |  |  |  |
| Parietal lobule | L | 943 | -36 | -66 | 50 | 5.05 |
| Parietal lobule | R | 276 | 40 | -52 | -28 | 5.18 |
| BA46 | L | 42 | -46 | 28 | 24 | 4.23 |
| Middle frontal gyrus | R | 50 | 34 | 2 | 58 | 4.10 |
| Middle frontal gyrus | L | 46 | -30 | -2 | 66 | 3.97 |
| SMA | L | 57 | -2 | 14 | 48 | 4.10 |
| BA10 | L | 30 | -42 | 44 | 0 | 4.10 |
| Substitution (different-same reference > different-same comparison) |  |  |  |  |  |  |
| SMA | L | 259 | -2 | 8 | 56 | 6.06 |
| Middle frontal gyrus | L | 83 | -50 | -44 | 50 | 5.65 |
| IPL | L | 37 | -44 | 2 | 50 | 6.53 |
| Updating (repeat reference > repeat comparison) |  |  |  |  |  |  |
| IPL | L | 302 | -48 | -42 | 50 | 6.17 |
| IPL | R | 82 | 34 | -44 | 38 | 5.61 |
| MFG | R | 49 | 38 | 4 | 58 | 5.54 |
| SPL | L | 22 | -34 | -64 | 52 | 5.42 |
| MFG | L | 10 | -28 | 0 | 66 | 5.21 |

**Figure 3:**
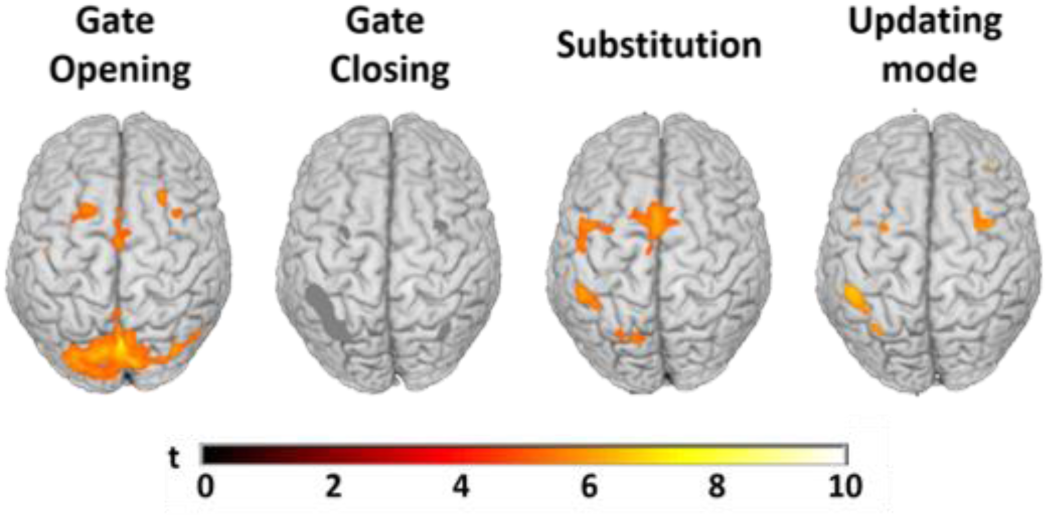
Whole-brain group search results for neural substrates of the different WM updating sub-processes/states, displayed as rendered dorsal 3D views (voxel-wise FWE *p* < 0.05, *K*_*E*_ > 10). The grey clusters presented in “gate closing” contrast refer to the clusters found using voxel-wise FDR threshold correction (*p* < .05)

The analysis of the gate-closing process (required on comparison trials that follow reference trials) did not yield significant activations with the conservative FWE whole-brain correction. To probe further for potential neural substrates of gate-closing, we applied a more lenient form of whole-brain correction (voxel-wise FDR *p* < 0.05, *K*_*E*_ > 10) to this contrast, which revealed primarily activity in bilateral PPC, specifically in the superior parietal lobule (SPL)/intraparietal sulcus (IPS), along with smaller clusters of dorsal frontal activation (for full listing of active clusters, see Table 3). When WM did not only have to be updated but information in WM had to be replaced (*substitution*, required on reference trials where the current stimulus mismatched the WM referent), activity was enhanced in left dlPFC (middle frontal gyrus) and inferior parietal lobule (for full listing of active clusters, see Table 3). Finally, being in an *Updating mode* was also associated with increased activation of dorsal frontal and parietal regions, including most prominently the left PPC.

In sum, in line with expectations, the exploratory whole-brain analysis identified core components of the FPN as supporting the regulation of WM updating/protection processes. However, these contrasts also suggest some regional differences, suggesting a relatively greater involvement of medial posterior parietal (and visual) cortex in gate opening, of more lateral posterior parietal regions in gate closing operations, and relatively stronger prefrontal involvement in the substitution process.

#### ROI analysis

Activity related to the different WM updating operations in *a priori* ROIs was examined with ROI-wide FDR correction, using a voxelwise threshold of FDR *p* < 0.05. Dorsal views/axial slices illustrating key findings are shown in Figure 4. A list of peak coordinates is shown in Table 4. Additionally, we extracted and analyzed mean activity estimates from each of the ROIs. While this analysis is necessarily less sensitive, as it averages activity over entire anatomical regions, it allowed us to further quantify the potential regional functional specializations with respect to WM updating sub-processes, and to ensure that the inferences derived from the ROI-based search do not simply reflect (quantitative) thresholding effects, but genuinely (qualitatively) different activity patterns. Figures 5 and 6 present mean beta values extracted for each ROI (broken down into nuclei, in the case of the BG). A summary of the entire statistical analysis, including Bayes factors, is presented in Table 5.

**Table 4:**
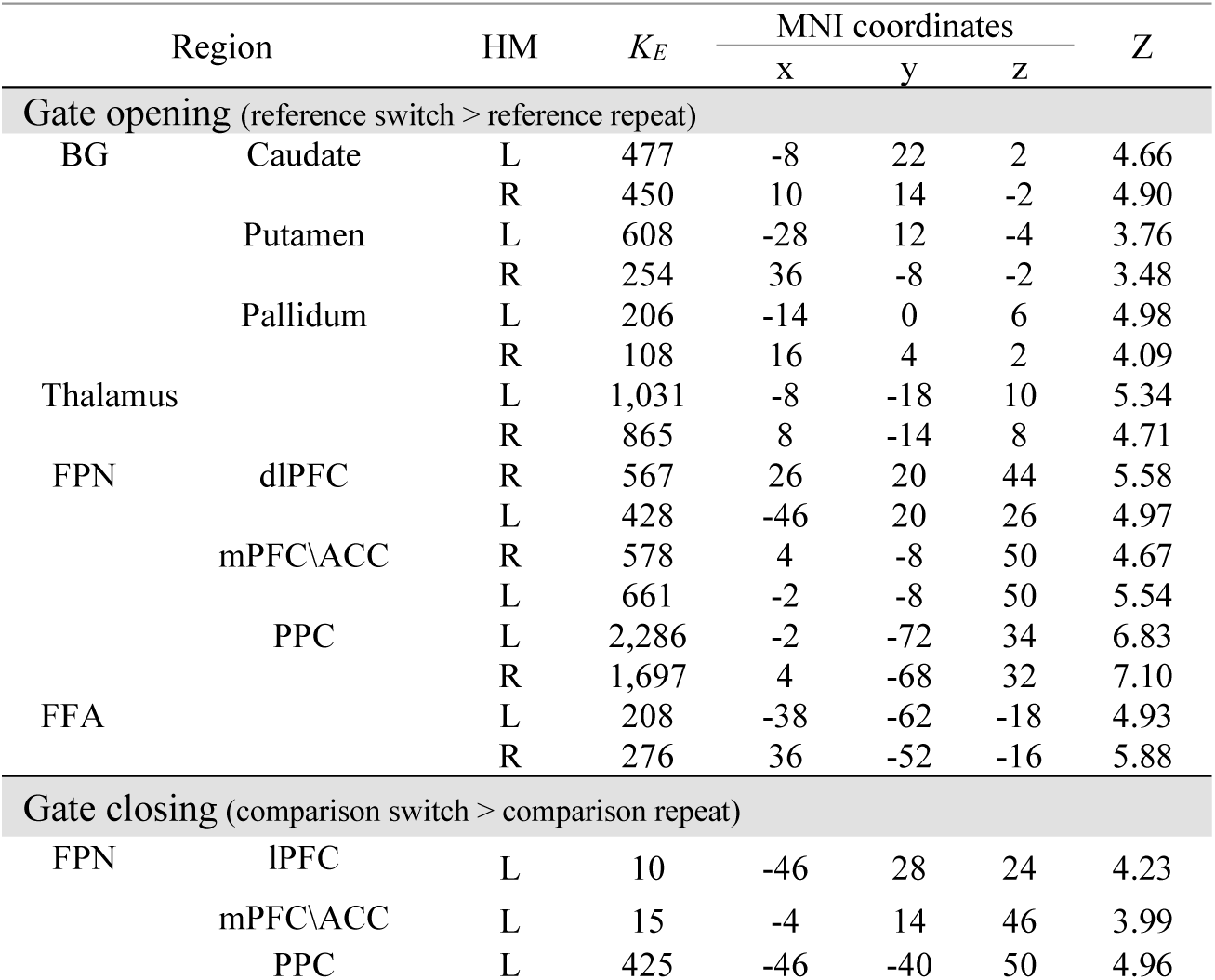
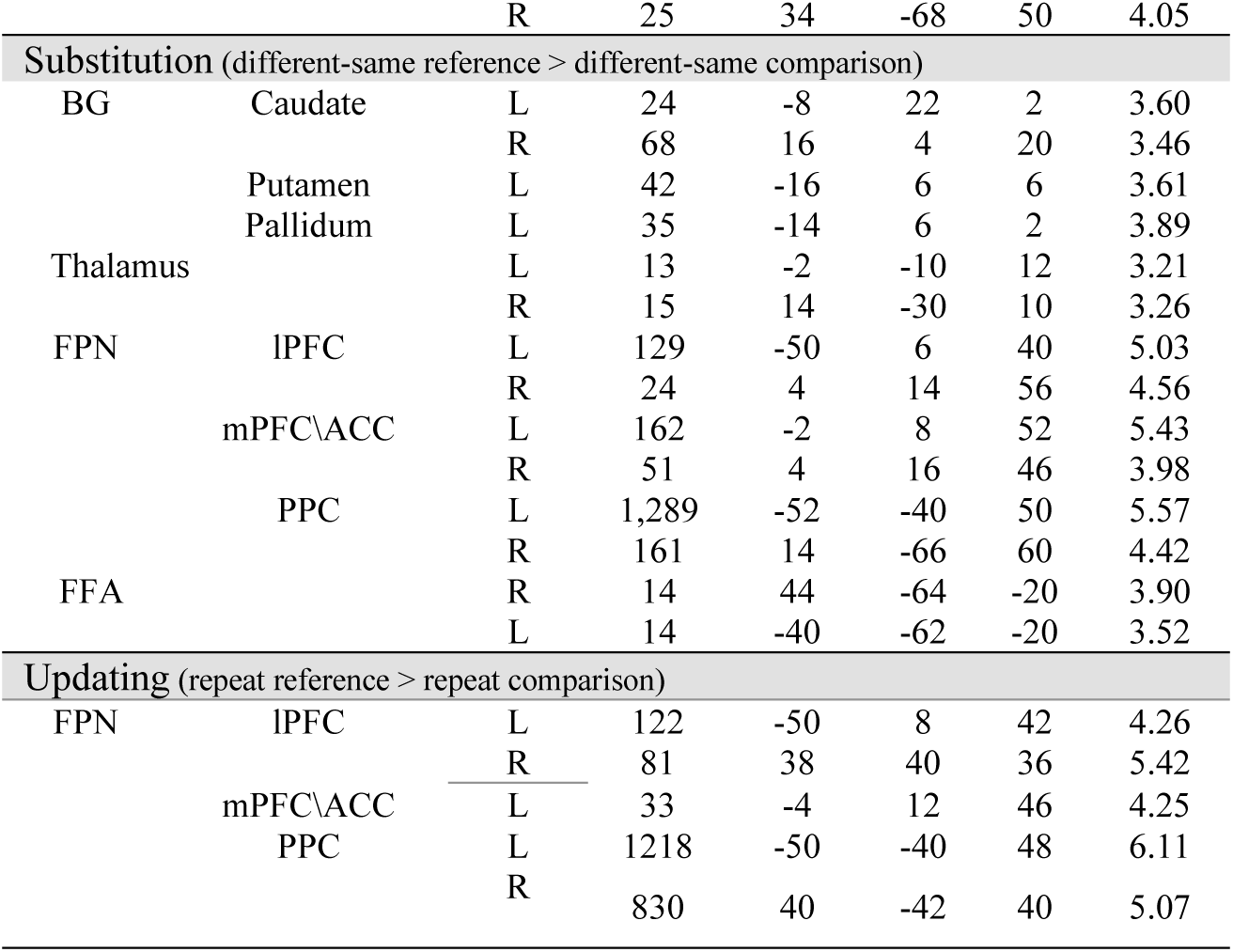
ROI analysis peak activation in MNI coordinates; Z refer to z-score at peak activated voxel

**Table 5:**
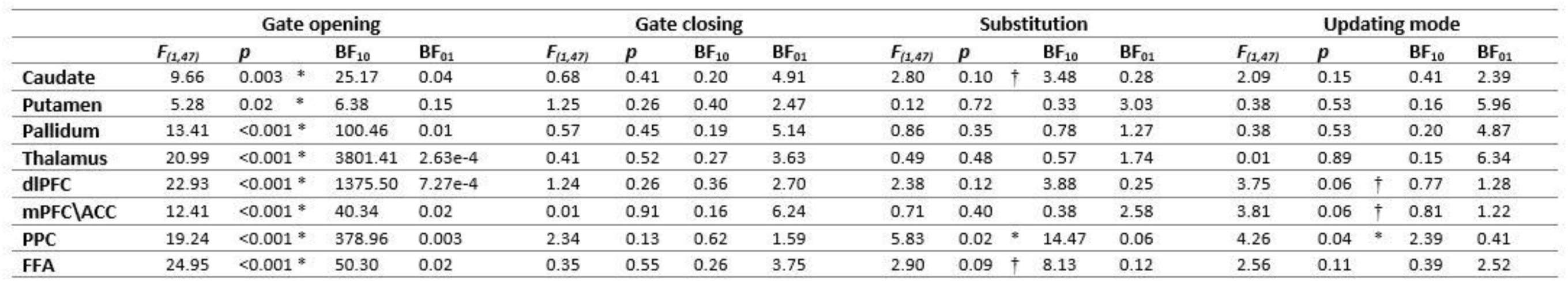
Summary of the statistical analysis on the mean activity beta-values, divided by ROIs and by the sub-processes. For each region X sub-process, a Bayes factors favoring the hypothesis (BF10) and the null hypothesis (BF01) were calculated as well. († denotes p < .1; * denotes p < .05).

**Figure 4:**
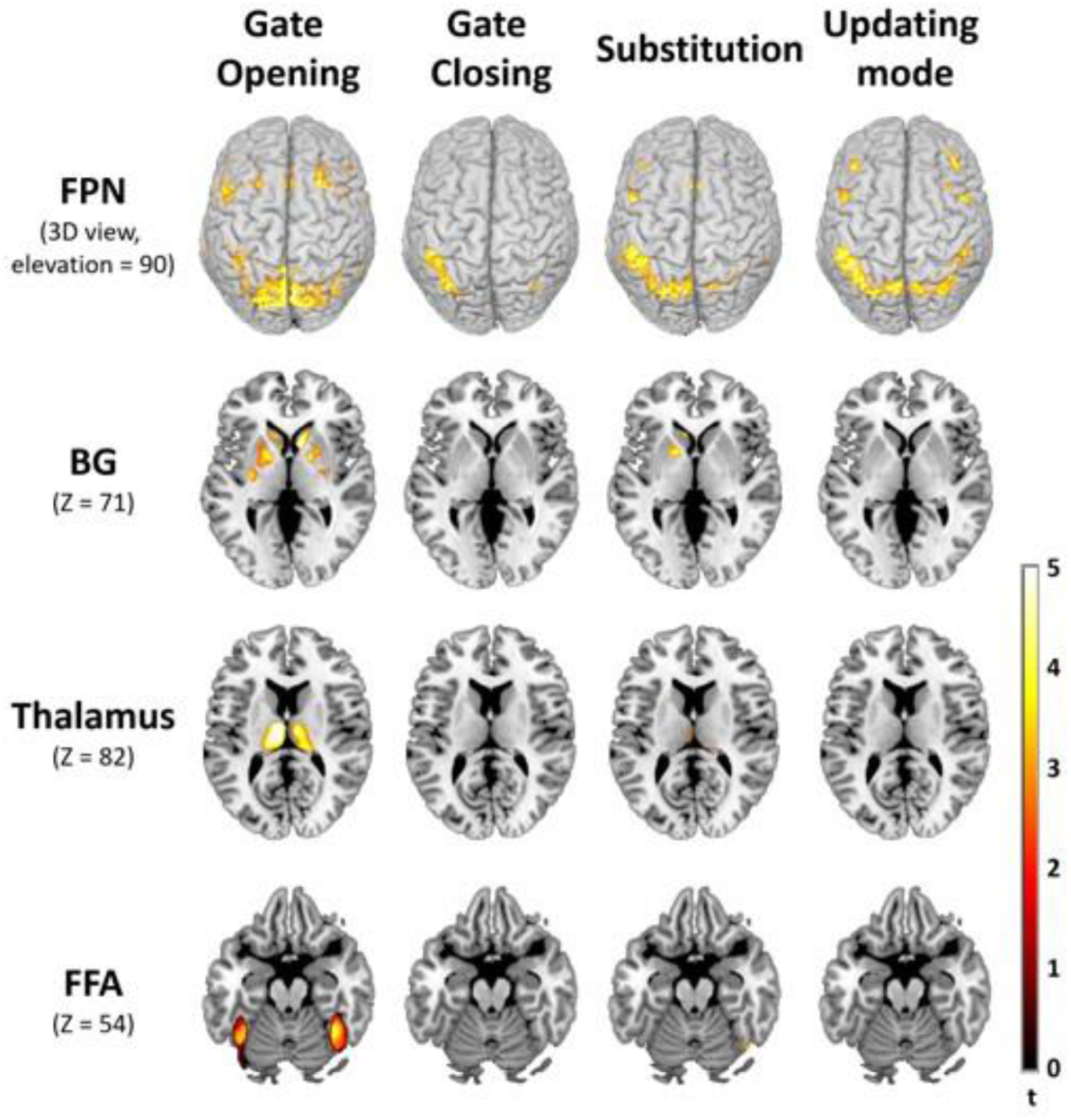
Results of the ROI analyses are displayed as a function of WM updating sub-process (columns) and ROI (rows) on dorsal-view rendered 3D brains (Top row) and axial slices (other rows). FPN = frontoparietal network, BG = basal ganglia, FFA = fusiform face area.

**Figure 5:**
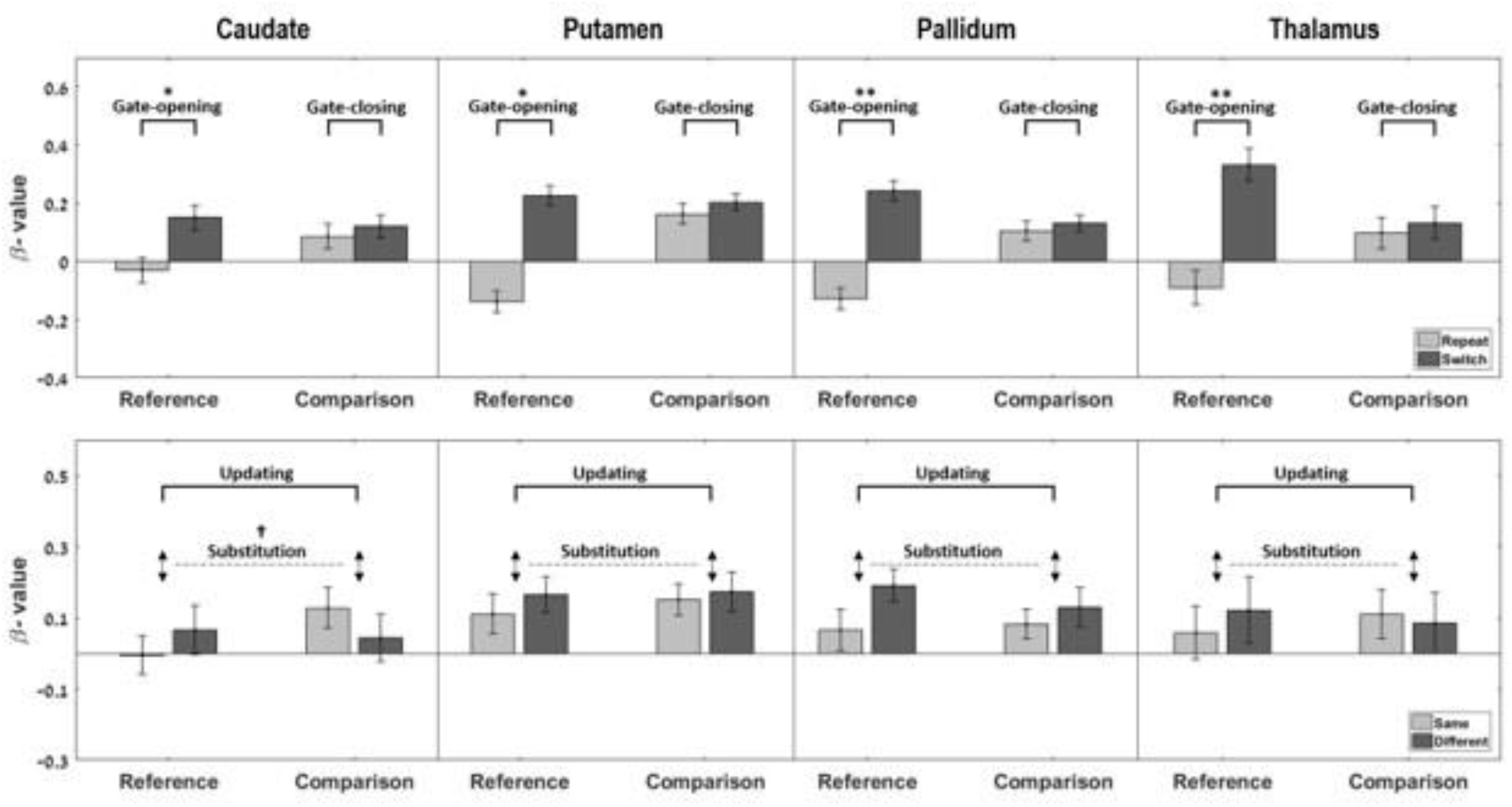
Mean activity estimates (and standard error of the mean) for the BG nuclei and thalamus are shown for reference and comparison trials as a function of whether the condition was repeated or switched to (upper panel) and of whether the stimulus/response was the same or different to the reference stimulus held in WM (lower panel). The figure also highlights the four key contrasts defining gate-opening, gate-closing, substitution and updating mode († denotes *p* < .01; * denotes *p* < .05; ** denotes *p* < .001).

**Figure 6:**
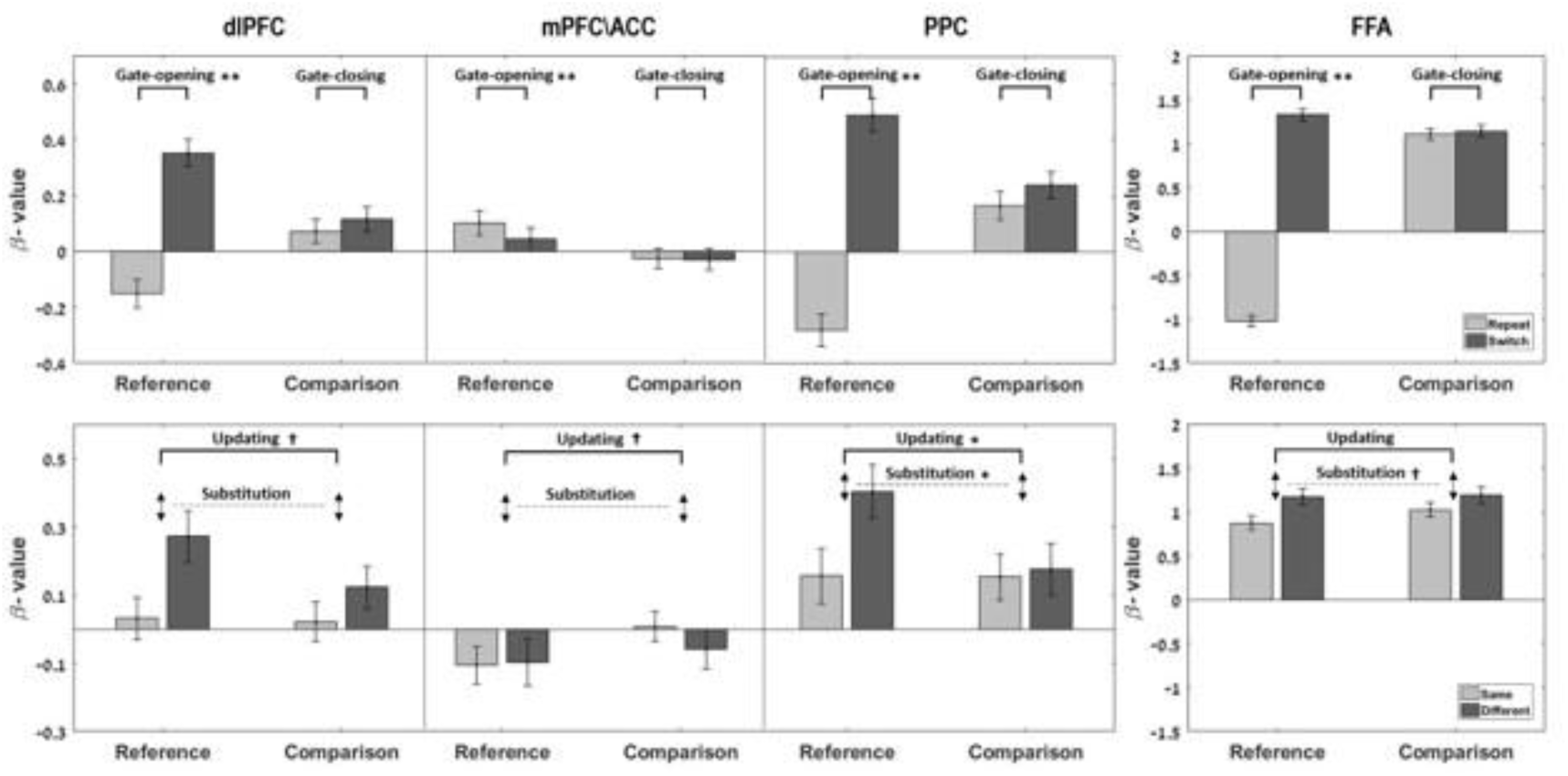
Mean activity estimates (and standard error of the mean) for the FPN components and the FFA are shown for reference and comparison trials as a function of whether the condition was repeated or switched to (upper panel) and of whether the stimulus/response was the same or different to the reference stimulus held in WM (lower panel). The figure also highlights the four key contrasts defining gate-opening, gate-closing, substitution and updating mode († denotes *p* < .01; * denotes *p* < .05; ** denotes *p* < .001).

### BG

In the ROI-based search, we observed enhanced activity in BG nuclei for gate opening and substitution, but not for gate closing or being in an updating mode (see Figures 4 and 5). More specifically, opening of the gate to WM was associated with the most widespread increase in activity, involving all of the BG nuclei bilaterally (Figure 5). In contrast, no activation increase was detected in any part of the BG during WM gate closing or updating mode. Finally, the substitution of old WM content with new information was associated with increased activation in the caudate, left putamen, and left pallidum.

This pattern of results was largely replicated in the analysis of mean activation estimates for individual nuclei (Table 5; Figure 5). Mean activation of the BG was significantly higher during switch than repeat trials in reference trials, but not in comparison trials. Bayes factors corroborated the implication of the BG in gate-opening processes, especially in the caudate and pallidum (*gate-opening*: BF_10_ caudate = 25.17; BF_10_ putamen = 6.38; BF_10_ pallidum = 100.46. *gate-closing*: BF_10_ caudate =0.20; BF_10_ putamen = 0.40; BF_10_ pallidum =0.19), and they also provided support against an involvement of these two nuclei in gate-closing (BF_01_ caudate = 4.91; BF_01_ pallidum = 5.14). Updating mode related activity was not significant for any contrast, and the non-involvement of the BG in that mode was supported by the Bayes factors results, which favored the null hypothesis (BF_01_ putamen = 5.96; BF_01_ pallidum = 4.87). Finally, only mean activity in the caudate also exhibited some evidence for an involvement in the substitution process (BF_10_ = 3.48). In sum, the present data support the longstanding proposal of BG involvement in input-gating of WM content, by showing that the BG are robustly associated with the process of opening the gate to WM. We also observed some evidence for substitution related activity, but – most importantly – supported by the Bayesian analysis results, the BG nuclei seem to play no active role in closing the gate to WM.

### Thalamus

Similar to the BG, the thalamus was found to display activity increases during WM gate opening, but displayed no detectable increase in activity during the gate closing operation or with respect to being in an updating mode. This was born out both by the search for significant clusters within the thalamus ROI (Figure 4), as well as by the analyses run on mean thalamus activation (Figure 5): Switch trials evoked higher mean activation than repeat trials in reference trials but not in comparison trials, and this gate-opening effect was strongly supported by Bayes factor analysis (BF_10_ = 3,801). By contrast, we observed some evidence against the thalamus’ involvement in gate-closing (BF_01_ = 3.63). Moreover, neither the substitution cost nor updating mode contrasts were significant, with the Bayes factor analysis speaking against thalamus involvement in the updating mode (BF_01_ = 6.34). In sum, as in the BG, we observed strong evidence for activity increase in the thalamus when the gate to WM had to be opened, whereas we observed some evidence against an involvement in gate closing.

### FPN

As already suggested by the whole-brain analysis above, we found that components of the FPN were activated by all WM updating sub-processes, but the pattern of activation was suggestive of a functional fractionation of the different FPN nodes (see Figures 4 and 6). Specifically, the search for significant activated clusters within the ROIs showed that, whereas the entire FPN was robustly and bilaterally activated during the gate opening operation and by being engaged in an updating mode, the process of WM content substitution produced much more lateralized activity in the left parietal and left lateral frontal cortex, and gate closing was associated almost exclusively with enhanced left parietal activation along the IPS (Figure 4).

The analysis of mean ROI activity and the Bayes factors confirmed this picture: As shown in Figure 6 (and in Table 5), the gate-opening contrast was significant for mean activation in all FPN components. Moreover, the Bayes factors favoring the hypothesis of these regions’ involvement in gate opening supported this pattern (BF_10_ dlPFC = 1,375.50; BF_10_ mPFC\ACC = 40.34; BF_10_ PPC = 378.96). In contrast, the gate closing contrast at the level of mean ROI activity was not found to be significant in any of the FPN components. Bayes factors favored the null hypothesis of no involvement in gate-closing for the mPFC (BF_01_ = 6.24) but did not support either hypotheses for the other components (see Table 5). The substitution cost contrast was significant for the PPC, with strong support from the Bayes factor analysis (BF_10_ = 14.47), but not the frontal FPN components, though Bayes factors indicated some evidence for the dlPFC’s involvement in substitution (BF_10 =_ 3.88). Finally, the updating mode contrast was significant for mean activity in the PPC and marginally significant for the frontal regions. The Bayes factors analysis did not provide support for or against this involvement, however (see Table 5).

In sum, in line with the results of the whole-brain analysis, the ROI-based analysis provided additional evidence for a functional dissociation of FPN components’ roles in WM updating, with the frontal and parietal nodes being concerned with gate opening and being in an updating mode, but posterior parietal cortex additionally contributing to substitution of information in WM. While the ROI-based search revealed some activity in the left PPC during gate closing, Bayesian analysis on the mean beta-values of PPC provided neither support for (BF_10 =_ 0.62) nor against (BF_01_ = 1.59) this region’s involvement in gate closing. Together, we view these data as merely suggestive of a possible involvement of PPC in gate closing.

### FFA

The (functionally defined) FFA ROI displayed a similar activity pattern to that observed in the BG, thalamus, and frontal (but not parietal) FPN components. In the search for active clusters, we observed loci in the FFA that showed significant activation increases during substitution, but the most pronounced activity was observed during gate opening (Figure 4). In contrast, no activity increase was detected during gate closing or in relation to the updating mode. The same pattern was confirmed in the analysis of mean FFA activation (Figure 7) and in the Bayesian analysis. Specifically, there was greater mean activation during switch trials than repeat trials in reference trials but not in comparison trials, and an involvement of the FFA in gate-opening was strongly supported by the Bayes factor analysis (BF_10_ = 50.30). Moreover, we found some evidence against the FFA’s involvement in gate closing, with the Bayes factor favoring the null hypothesis (BF_01_ = 3.75). Substitution cost and updating mode contrasts were not significant at the mean ROI activity level, however, the Bayes factor indicated evidence in favor of the FFA’s involvement in substitution (BF_10_ = 8.13). Thus, intriguingly, processes related to allowing sensory content to enter WM (gate opening) and/or to replace WM representations (substitution) seem to have a direct impact on activity in the sensory regions that are involved in processing and/or representing the relevant stimulus material.

## Discussion

There is copious evidence for the involvement of the FPN, BG, and thalamus in maintaining and updating WM content, but how the regions may differentially contribute to different sub-processes of WM updating is not well understood. To address this important question, we combined fMRI with the recently developed reference-back task, which enabled us – for the first time - to tease apart neural substrates of gate-opening, gate-closing, and substitution processes, as well as an updating mode of operation. Moreover, we used face stimuli in order to examine updating-related activity in sensory cortex specialized for processing the WM items, namely the FFA.

Our behavioral results fully replicated previous studies using this protocol (Kessler, 2017; Rac-Lubashevsky & Kessler, 2016a, 2016b). Moreover, the imaging data indicate that the FPN, BG, and thalamus all contribute to gate-opening and substitution processes. Being in an updating mode associated with FPN activity, mainly the PPC, but not the subcortical components. Also, we found that FFA activity displayed robust effects of WM gate opening and substitution processes.

### Gate opening, substitution, and updating mode in the WM network

The use of the reference-back paradigm allowed us to isolate different sub-processes involved in WM updating processes, thus enabling more precise process-to-brain region attribution and, accordingly, a stronger test of the PBWM model (Frank et al., 2001; O’Reilly & Frank, 2006). In general support of that model’s proposal of BG-thalamus-PFC circuits supporting WM, we observed involvement of all components of this network (as well as PPC) in the processes of opening the gate to WM, and in subsequently replacing the current content of WM with new perceptual information (substitution). The PBWM model posits that it is specifically the BG (and not the FPN) that are implementing the gating operation. However, under the assumptions of the model it is nevertheless plausible that one would also observe gate opening and substitution related activity more broadly throughout the WM network, as the opening of the gate, and in particular the substitution process, would be expected to have knock-on effects in the thalamus and FPN. Moreover, in support of the model’s differentiation between the gating mechanism in the BG and the representation of WM content in FPN, we found that an “updating mode”, which refers to an open-gate state that does not involve any change in gating status, was associated exclusively with FPN, and not with BG or thalamic activity. Hence, it appears that opening the gate to WM relies on activating the fronto-thalamic-striatal loop, but only frontoparietal cortex is involved in keeping the gate in an open state to allow for continuous updating.

### Gate-opening versus gate-closing

Intriguingly, whereas we observed strong evidence for the BG, thalamus, and FPN regions’ involvement in gate-opening, we did not observe strong evidence for any of these region’s involvement in gate closing, but substantial evidence *against* such an involvement for the BG and thalamus. Previous behavioral studies demonstrated that both opening and closing the gate to WM involves a reaction time cost. This was observed both using the reference-bask task (Rac-Lubashevsky et al., 2017; Rac-Lubashevsky & Kessler, 2016a, 2016b, 2018) and in a sequence updating task (Kessler & Oberauer, 2014, 2015). Here we find that, under roughly equivalent behavioral costs of opening and closing the gate, the neural mechanisms involved in the two operations appear to be distinct. Notably, supported by Bayesian analysis, the BG and the thalamus showed clear single dissociations – with strong evidence for gate-opening related activity, and strong evidence against gate closing related activity. However, we did not observe the obverse pattern in any other region. In fact, the only region where we detected any sign of potential involvement in gate closing was the PPC, but the evidence was not conclusive.

We offer three possible interpretations of these results that raise interesting questions for follow-up studies. First, the lack of a BG activation increase in relation to gate closing *per se* may plausibly reflect the fact that the closed gate (i.e., tonic inhibition of the thalamus) represents a BG default state (e.g., Chevalier and Deniau, 1990), returning to which might not impose additional local metabolic demands on the BG. In other words, the observation of robust behavioral gate closing costs (both in the current and prior studies), combined with the lack of strong evidence for any one region’s contribution to closing the gate to WM, suggests that gate closing may be a time-consuming but not an active process. This possibility fits with the PBWM model suggesting that the closed gate represents a default state. Second, based on our observation of some (though not strong) evidence for an involvement of the PPC in gate closing, it is also possible that gate closing does require active cortical engagement, and that this process originates in the PPC. The PPC has been frequently implicated in WM maintenance (e.g., Majerus et al., 2016; Quentin et al., 2019), and it has ample direct anatomical projections to the BG (e.g., Cavada and Goldman-Rakic, 1991; Jarbo and Verstynen, 2015), thus making it a plausible contributor to gating processes. However, given the inconclusive nature of its involvement in gate closing in the present study, future studies, perhaps involving targeted neuro-stimulation of the PPC, are needed to rigorously test this possibility.

Third, another possible account for the lack of active BG (and thalamic and FPN) involvement in the gate closing process could be that closing the gate to WM relies on direct projections from the VTA to frontal regions, and hence does not activate the BG (cf. Braver & Cohen, 2000; Chatham & Badre, 2015; D’Ardenne et al., 2012). Specifically, while gate opening should be selective by nature, and hence involve activating specific BG-PFC loops, gate closing may be non-selective and hence could operate through the (non-specific) VTA-PFC pathway, and our imaging protocol may not have been sensitive enough to detect metabolic changes in the VTA, being very small midbrain region. Future studies could potentially probe whether a different neural signature emerges under conditions where gate-closing has to be selective, too, such as when items from multiple categories have to be updated independently, and/or employ fMRI protocols optimized for gauging VTA activity.

Finally, it should be noted that although in our design gate switching always involved a change in the color of the frame (from red to blue or vice versa), the dissociation between gate opening and closing, and the brain regions involved in the two, makes it extremely unlikely that the gating effects merely reflect the perceptual effects of switching between frame colors. In such a case, symmetrical effects of gate opening and gate closing were expected, unlike our clear indications for a dissociation between the two. Moreover, while it is tempting to interpret gate-switch costs (opening and closing) as task switch costs, several empirical and theoretical considerations argue against this interpretation. On a theoretical level, such an interpretation suggests that being in an open-gate state represents a different task set than being in a closed-gate state. However, what this view would consider to be switching between two “tasks” is identical to what we refer to as switching between the gate states. Hence, appealing to the notion of task switching would not represent an alternative account but a re-labeling of the gate switching operations, and one that in our view is theoretically less coherent. Empirically, the interpretation of gate switching as task switching implies similar neural correlates for gate-opening and gate-closing contrasts (as both compare “task switches” to “task repetitions”). In contrast to this prediction, we observed distinct activity for the two, including a conjunction analysis that did not yield any overlapping brain areas between gate opening and gate closing. Finally, previous behavioral work found evidence for task-switching and gate-opening operations to proceed in parallel, thus implying that they are distinct processes (Kessler, 2017).

### The role of posterior cortex in WM updating

Many prior studies have established that the PFC is a crucial component of short-term memory (Fuster & Alexander, 1971), cognitive control (Miller & Cohen, 2001) and WM updating and maintenance (e.g., Narayanan et al., 2005). However, the specific role it plays in supporting these abilities is still under debate. One traditional view, which is also implemented in PBWM, holds that the PFC plays a key role in encoding and storing goal-directed representation within WM (Goldman-Rakic et al., 1996; Courtney et al., 1998; O’Reilly and Frank, 2006). In contrast, a more recent view (the *sensory recruitment hypotheses*) does not identify the PFC with temporary storage *per se*, but rather characterizes the lateral PFC as responsible for directing selective attention towards memory representations, with the actual representations being held in the posterior cortex, that is, in areas that also serve the perception and long-term storage of the memoranda (D’Esposito et al., 2000; Feredoes et al., 2011; Lara & Wallis, 2015; Postle, 2006; Serences, 2016).

Relevant to this debate, our findings demonstrate the involvement of the FFA in WM updating sub-processes. In particular, gate opening and substitution processes, but not gate-closing, were associated with elevated neural activity in the FFA. This finding supports the notion that posterior, “perceptual” regions play a major role not only in WM maintenance (as already posited by the sensory recruitment account), but also in WM updating. We offer two tentative accounts for this finding. The first is that the posterior cortex is more heavily recruited for WM maintenance in situations that call for substitution of the information, or for switching from “passive maintenance” (as reflected in comparison trials) to updating. A second, not mutually exclusive interpretation is that the same signaling cascade that leads to gating information into WM also serves to concurrently boost attention to the to-be-encoded items represented in posterior perceptual regions. While input gating involves a feed-forward flow of information from perception to WM, representations held in WM may in turn lead to directing attention toward perceptual regions, in a concurrent feed-backward fashion. These possibilities could plausibly be evaluated by combining the reference-back task with more time-sensitive measures of neural activity in future studies.

## Conclusions

To conclude, the present study revealed for the first time distinct neural activity related to gate opening, gate closing, substitution, and updating mode, by combining the reference-back protocol with fMRI. While gate opening was associated with activation of the BG-thalamus-PFC loop, in accordance with the PBWM model, the same was true for FFA activity. Moreover, we observed strong evidence against an active involvement of the BG and thalamus in the gate closing process. These results supply important novel data to inform our evolving theories of the neuroanatomical mechanisms supporting working memory.

## Acknowledgments

This study was funded by US-Israel Binational Science Foundation (BSF) grant awarded to YK and TE.

## Notes

**Conflict of interest statement:** The authors declare no competing financial interests.

### Competing Interest Statement

The authors have declared no competing interest.

### Summary of Updates

The section on the paradigm updated to clarify issues of removal and repetition suppression. Also, to elaborate on mapping the paradigm contrasts onto the PBWM model. Figure 1 revised. The discussion updated to clarify issues of task-switching and a minor elaboration on the interpretations of the gate-closing results.

